# High-throughput phenotyping of leaf discs infected with grapevine downy mildew using shallow convolutional neural networks

**DOI:** 10.1101/2021.08.19.456931

**Authors:** Daniel Zendler, Nagarjun Malagol, Anna Schwandner, Reinhard Töpfer, Ludger Hausmann, Eva Zyprian

**Affiliations:** Julius Kuehn Institute, Institute for Grapevine Breeding Geilweilerhof, Siebeldingen, Germany

**Keywords:** Downy mildew, *Vitis vinifera*, CNN, high-throughput, phenotyping, leaf discs

## Abstract

Objective and standardized recording of disease severity in mapping crosses and breeding lines is a crucial step in characterizing resistance traits utilized in breeding programs and to conduct QTL or GWAS studies. Here we report a system for automated high-throughput scoring of disease severity on inoculated leaf discs. As proof of concept, we used leaf discs inoculated with *Plasmopara viticola* causing grapevine downy mildew (DM). This oomycete is one of the major grapevine pathogens and has the potential to reduce grape yield dramatically if environmental conditions are favorable. Breeding of DM resistant grapevine cultivars is an approach for a novel and more sustainable viticulture. This involves the evaluation of several thousand inoculated leaf discs from mapping crosses and breeding lines every year. Therefore, we trained a shallow convolutional neural-network (SCNN) for efficient detection of leaf disc segments showing *P. viticola* sporangiophores. We could illustrate a high and significant correlation with manually scored disease severity used as ground truth data for evaluation of the SCNN performance. Combined with an automated imaging system, this leaf disc-scoring pipeline has the potential to reduce the amount of time during leaf disc phenotyping considerably. The pipeline with all necessary documentation for adaptation to other pathogens is freely available.

## 1. Introduction

Objective and standardized analysis of phenotypic traits is one of the most crucial requirements of modern plant breeding and genetic mapping programs. Association studies like quantitative trait locus (QTL) analysis and genome wide association studies (GWAS) are highly dependent on precise genotypic and phenotypic data. However, in breeding research it is still common to rely on visual assignment of categorical classes describing for example disease severity caused by certain pathogens. To reduce the possibility of human errors, it is advisable that these class assignments are performed by multiple people on multiple biological replicates of the genetic material under study and in multiple repetitions of the tests. Further, in the case of host/pathogen interactions it is sometimes important to screen material with different isolates or races of the pathogen at different stages of the infection to identify the durability or resistance spectrum of a novel resistance source. These considerations make obvious, that manual phenotyping is a tedious and time-consuming task when done properly. In addition, albeit a high degree of experience of personnel performing the scorings, subjectivity of individual persons always introduces a certain degree of bias.

This is where computer vision comes into play. The advancements in this field during the last decades have made it possible to automatically collect large numbers of high-quality images to record and store different stages e.g. during infection progression. Moreover, machine learning approaches have been developed over the last one and half decades that exceed former technology. Machine learning is now available in all sorts of technologies allowing e.g. speech recognition, face recognition or pattern recognition in written text, and more applications. Additionally, the accessibility of machine learning algorithms has been greatly facilitated when the free and open source library TensorFlow was made available by Google in 2015 [1,2]. For image classification, deep convolutional neural networks (DCNN) have been developed starting from Alexnet [3] going to VGG [4] over to Inception and GoogLeNet [5], always increasing the complexity and the number of layers making them even “deeper”. These state of the art DCNNs have an excellent accuracy when trained properly. However, they require comprehensive computational resources. In recent years more and more articles were published using such networks in various combinations and adaptations to identify and quantify diseases on a wide variety of plants under various scenarios (Review articles on these topics [6], [7]). The first attempt of implicating deep learning for image-based plant disease assessment carried out in 2016 successfully classified 14 different crops and 26 diseases with an average accuracy of 99% [8]. A recent study combines several of these DCNNs for detecting and to distinguish different grapevine diseases on whole leaves [9]. In a further study deep learning approaches using CNNs were employed to detect grapevine downy mildew and spider mites under field conditions on images of whole plants [10].

In this study we decided to use so-called “shallow convolutional neural networks” (SCNNs) which use only a small number of image convolutions and subsequent fully-connected layers thereby reducing the requirement of high computational resources and training time [11,12]. As proof of concept, the SCNNs were trained to detect DM infections on RGB images of experimentally inoculated leaf discs.

DM caused by *P. viticola [(Berk. & Curt*.*) Berl. & de Toni]* is one of the major diseases’ viticulture is facing annually in temperate climate areas [13,14]. In 2016 and 2021 very heavy epidemics occurred in German viticulture with a strong economic impact due to intensified plant protection regimes and yield losses. The obligate biotrophic oomycete invades green plant tissue via zoospores, which emerge from vegetative generated sporangia or sexually produced oospores during wet and humid conditions. Using their flagella, zoospores swim to stomata, encyst at their rim and form a primary hypha, which enters the substomatal cavity, branches between spongy mesophyll cells and forms haustoria. After three to four days under optimal conditions, the oomycete forms so-called sporangiophores that emerge through stomata. Sporangia are formed at their tips and finally dispersed to restart the infection cycle under humid and wet conditions [15]. DM can have a devastating impact on yield when young inflorescences up to pea-sized berries are infected. The cultivated grapevine *Vitis vinifera* ssp. *vinifera* which is grown world-wide has no natural resistance against this pathogen. Grapevine growers therefore rely on strict plant protection regimes that require high amounts of fungicides [13,14]. To reduce the amount of plant protection chemicals used in viticulture, breeding programs have been established that focus on the introgression of natural resistances against this and many other pathogens into new breeding lines, eventually creating new resistant cultivars for future sustainable viticulture [16–18].

These breeding efforts first require screening of segregating crosses for the identification of novel resistance traits, which then, in the next step, can be genetically identified and introgressed into existing elite lines. For experimental inoculations of DM under controlled environmental conditions, leaf disc assays are preferably used and the degree of disease severity determined by visual assessment [19–23]. The amount of leaf discs for these screenings easily reaches several thousand per year making the need for a high-throughput screening method obvious.

Following the approach used by Bierman et al. [24] where they used a DCNN for quantification of powdery mildew infections on leaf discs, we trained SCNNs to automatically detect areas on leaf discs with DM infections using small number of images. To ensure accuracy of the trained SCNNs we created ground truth data and calculated the true positives. After observing a true positive ratio of 95 % in average, we implemented an easy to use leaf-disc-scoring pipeline that can be run on any given Unix system. The pipeline is available as an open-source GitHub repository with instructions included on how to train SCNNs for other traits or pathogen symptoms https://www.github.com/Daniel-Ze/Leaf-disc-scoring.

## 2. Materials and Methods

### 2.1 Plant material

Infection experiments and image capturing was carried out on F_1_ individuals of two cross populations with different genetic backgrounds. The first population consisted of 497 F_1_ individuals of a cross between ‘Morio Muskat’ (‘Silvaner grün’ x ‘Muscat à petits grains blancs’) and a genotype COxGT2 (*V. coignetiae* x ‘Gewürztraminer’). The second population was derived by a cross between ‘Cabernet Dorsa’ (‘Blaufränkisch’ x ‘Dornfelder’) and Couderc 13 (*Vitis aestivalis* var. *lincecumii* x Couderc 162-5) and comprises 314 F_1_ individuals. The breeding line COxGT2 (Nursery E. Schrank, Dackenheim, Germany) and the cultivar Couderc 13 were described to exhibit a high degree of field resistance against downy mildew. Both crosses were performed in 2018 and seeds germinated in spring of 2019. All individuals were grown as potted plants in the greenhouse at the Institute for Grapevine Breeding Geilweilerhof in Siebeldingen (49°13′05.0″ N 8°02′45.0″ E) in the southwest of Germany with a single shoot of approx. 40 cm height and a pot diameter of 16 cm. The plantation was protected against powdery mildew until one week before an artificial leaf disc inoculation experiment.

### 2.2 Inoculation experiments

Leaf disc inoculation assays were performed in two and three independent replicates, respectively, for both cross populations during summer of 2020 to determine the levels of leaf resistance to downy mildew in the F_1_ progenies. Two leaves of each individual were sampled from the third and fourth apical node respectively and four leaf discs (one cm diameter) were punched using a cork borer. Leaf discs were arrayed upside-down on 1% water agar (Insula Agar Agar Pulver E406, Gustav Essig GmbH & Co. KG, Mannheim, Germany) in 245 mm Square BioAssay Dishes (Corning®, Corning, New York, USA). They were laid out according to a template of two 96 sample grids as shown in Figure 1. The template was designed to fit the dimensions of a Zeiss AxioZoom v16 motorized table.

**Figure 1:**
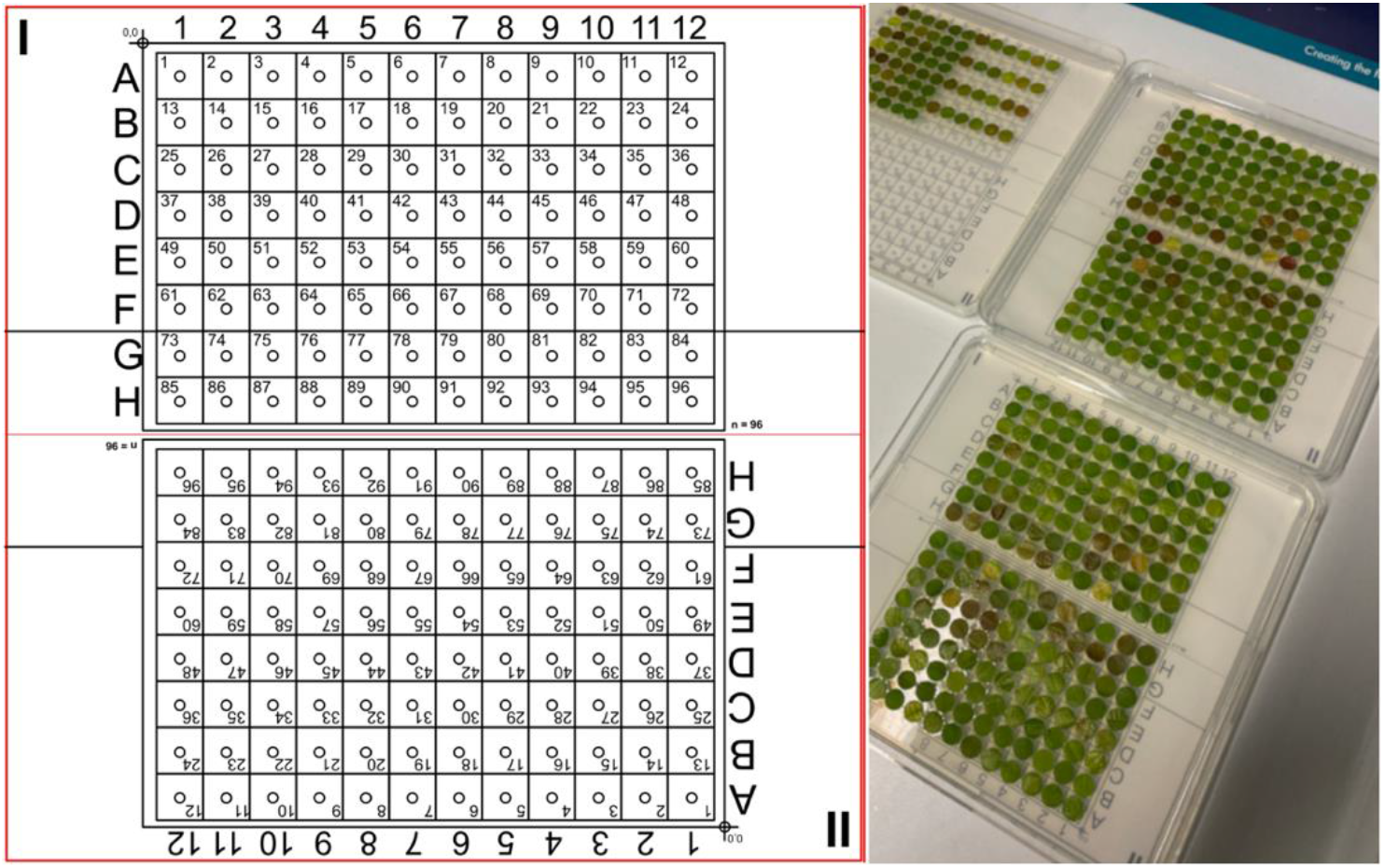
Template used for laying out leaf discs on a square 24.5 x 24.5 cm Petri dish filled with water agar and a picture of square dishes with leaf discs on them.

Three of the four leaf discs were artificially inoculated with a centered drop (30 µL) of a *P. viticola* sporangial suspension (18,000 sporangia per mL), the fourth leaf disc served as a water inoculation control. Sporangia were collected from leaves of un-sprayed susceptible *V. vinifera* vines from the field at the Julius Kühn Institute, Institute for Grapevine Breeding Geilweilerhof, Germany by vacuum suction using a water jet pump. Fresh sporulation was obtained by incubation of leaves that showed characteristic oil spots over-night under dark and wet conditions. After an overnight incubation at room temperature in the dark, spore suspension drops were removed using a water jet pump. Infected leaf discs were incubated for four days at 22°C with a photoperiod of 16 h (full spectrum light) and high relative humidity within the dishes.

### 2.3 Automated imaging of leaf discs

For automated imaging of leaf discs after inoculation with *P. viticola* a Zeiss AxioZoom V16 with motorized table and a 0.5X magnification lens was used (PlanApo Z 0.5x/0.125; Free Working Distance 114mm). Images were recorded in 10.5-fold magnification. A movement template was generated with the ZenBlue 3.0 software which allowed to automatically access the 96 positions. A sample template which can be directly loaded into Zen Blue 3.0 is available in the associated GitHub repository. The spacing of the template was calculated to be sufficient for leaf discs with one cm diameter. A LED ring light, two gooseneck lights and a backlight were used for illumination of the leaf discs. The gooseneck lights were positioned in an 45° angle to properly illuminate sporangiophores standing out from the leaf discs. The three lights combined allowed an exposure time of only 18 - 20 ms, making the whole imaging process fast. Consistent lighting of leaf discs was crucial for the trained CNN to reliably classify images.

Together with the motorized table, the Zeiss AxioZoom V16 is equipped with a software-autofocus, which was used to find the proper focal height for each leaf disc prior to imaging. To gain the highest focal depth, the smallest aperture possible was chosen. For each leaf disc one image was taken. In total 96 leaf discs could be imaged in roughly 15 min. A detailed protocol for the whole imaging process is available in the associated GitHub repository together with the complete workflow which can be directly imported into Zen Blue 3.0. With the workflow imported, imaging of leaf discs can be performed right away (https://github.com/Daniel-Ze/Leaf-disc-scoring/tree/main/microscope_instructions).

After imaging the first 96 samples, the plate was rotated 180° and the second half of the plate was recorded. Images were exported as JPEG files with a resolution of 2752 × 2208 pixels for further analysis. The exported images were renamed using a naming template and a Python script (available in the GitHub repository).

### 2.4 Manual phenotyping

Disease symptoms on leaf discs were captured at 4 dpi (days post-inoculation) using the AxioZoom V16 stereo microscope and were visually rated for disease severity. A reversed five-class OIV 452-1 descriptor (OIV, 2^nd^ edition 2001, https://www.oiv.int/en/technical-standards-and-documents/description-of-grape-varieties/oiv-descriptor-list-for-grape-varieties-and-vitis-species-2nd-edition) scale was used for the visual assessment. Disease severity was determined by the intensity of sporangia developed on the leaf discs. Where, score **1**: no sporangiophores (highly resistant), score **3**: one to five sporangiophores (resistant), score **5**: six to twenty sporangiophores (moderate), score **7**: more than twenty (susceptible), and score **9**: dense uniform cloud of sporangiophores (highly susceptible).

### 2.5 SCNN: training, eveluation and pipeline building

#### 2.5.1 Training and validation of the SCNNs

Leaf disc images representing the five possible inverse OIV452-1 classes were chosen from inoculation experiments with F_1_ individuals of the cross ‘Morio Muskat’ x COxGT2. The leaf discs were each sliced into 506 segments and manually classified into the classes background, leaf and leaf disc with sporangiophores (Figure 2) using image-sorter2 [25]. A script implemented in Python was used for random sampling of image slices from the respective classes for training and validation. The script is available in the associated GitHub repository.

**Figure 2:**
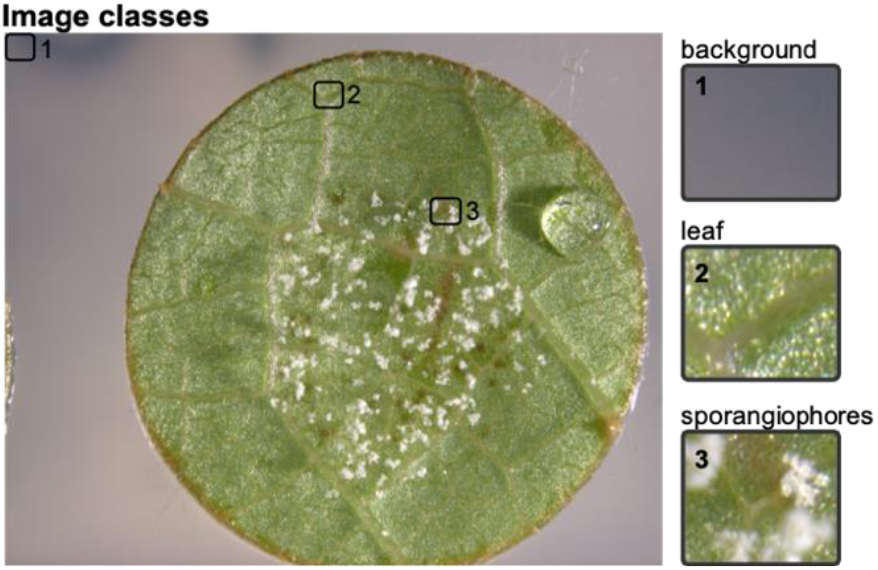
Example leaf disc picture with the three possible classes: 1 background = wateragar, 2 leaf = leaf disc partial image without infection, 3 sporangiophores = leaf disc partial image with sporangiophores.

The first binary SCNN has the classes “background” and “leaf disc” (CNN1). It was trained with 2919 images and validated with 947 images per class. The second binary SCNN was trained to classify images as “sporangiophores” and “no sporangiophores” (CNN2).

The final dataset comprised 968 images per class for training and 437 images per class for validation. All datasets are available in the GitHub repository. All SCNN trainings were performed on a desktop PC running Ubuntu 20.04, a 24 thread CPU and 32 Gb of RAM. Due to the use of SCNNs all calculations were performed using the CPU.

The detailed structure of the SCNN layers can be found in the GitHub repository. To retrain the SCNNs for special applications the scripts can be used with Tensorflow (v2.3.1) and Keras (v2.4.3) (GitHub repository) [2,26]. For the generated CNN1 model we used three image convolutions and one fully connected dense layer with 512 nodes (Figure 3, CNN1). As optimizer, “SGD” with “nestrov momentum” with a learning rate of 0.1 and a dropout of 0.4 was chosen (GitHub repository, CNN1.py). The structure of CNN2 was different. An additional convolution layer and two dense layers with 512 and 1024 nodes respectively were added (Figure 3, CNN2). Best results for the CNN2 were achieved with the “Nadam” optimizer with a learning rate of 0.0001 and a dropout of 0.5 (GitHub repository, CCN2.py). For training of the two SCNNs 30 epochs each were enough to achieve sufficient accuracy. Final image datasets for training the two SCNNs are available for download together with a Jupyter notebook containing detailed instructions (https://github.com/Daniel-Ze/Leaf-disc-scoring/tree/main/scripts).

**Figure 3:**
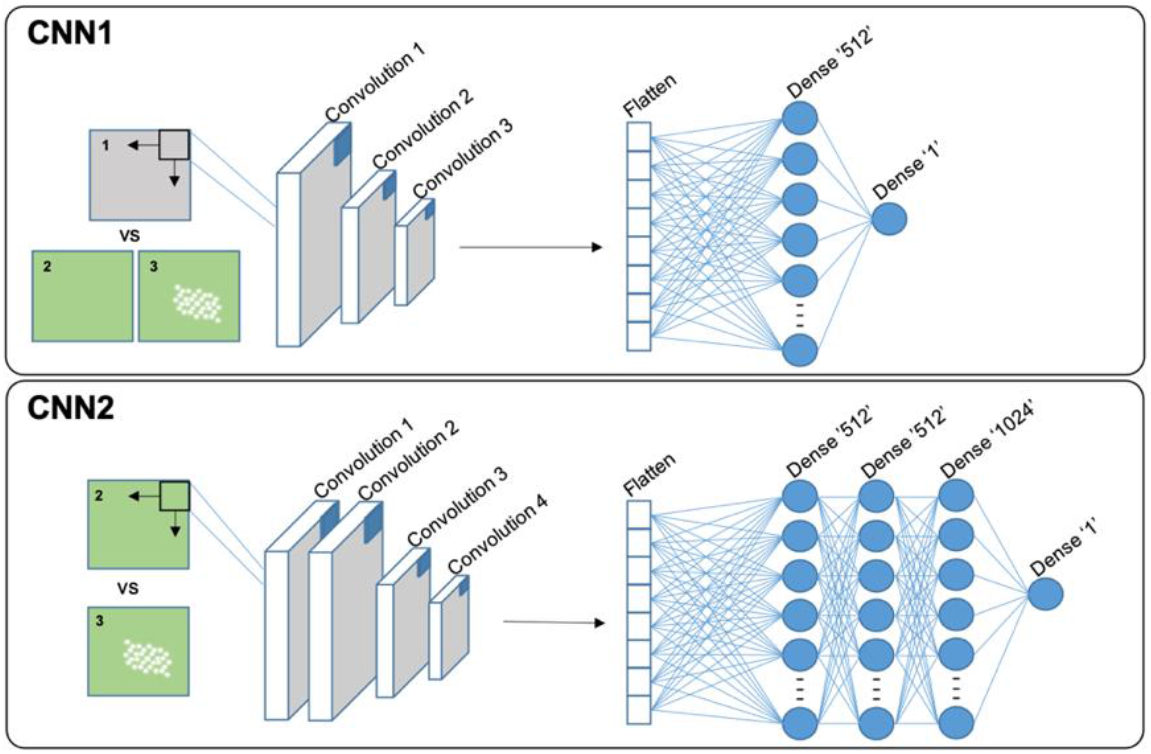
Structure of the two SCNNs. Each convolution layer consists of: convolution (Cov2D), activation (ReLU = rectified linear unit activation function) and pooling (MaxPooling2D). The fully connected layers (“Dense”) also have the ReLU activation. The last “Dense” ‘1’ layer has the Sigmoid activation function required for the binary output of the CNNs. The number of nodes in each “Dense” layer are indicated and further details can be found in the GitHub repository.

#### 2.5.2 Evaluating the performance of the trained SCNNs

To confirm the accuracy of the trained neural network, ground truth data was generated. Three independent experts classified slices of 30 leaf discs manually. For this test two different genetic backgrounds were used. First, 15 leaf disc images were chosen from the ‘Morio Muskat’ x COxGT2 population (three of each inverse OIV452-1 phenotypic class). The two SCNNs were originally trained with leaf disc images from this cross. To test the performance of the two SCNNs additional 15 images from a second independent cross (‘Cabernet Dorsa’ x Couderc 13) were analyzed (three of each inverse OIV class).

In total, 15180 image slices were sorted into the three different classes (“background” = 1, “sporangiophores” = 2, “no sporangiophores” = 3). It was ensured that a set of leaf discs was chosen which has not been used in the earlier training of the SCNNs. From this manually assigned set of scores, the true positive percentage was calculated by comparing the manually assigned classes of three independent experts with the classes of the two SCNNs.

Further, the neural network calculated percentage leaf disc area with sporangiophores was correlated with either the recorded inverse OIV452-1 score or the manually scored percentage leaf disc area with sporangiophores. The data was correlated using the R cor() package with the method “spearman” (OIV) and “pearson”, respectively (percentage leaf disc area with sporangiophores) [27].

#### 2.5.3 The classification pipeline

To make the analysis of a large set of images easier a simple one-line command line pipeline was implemented (Figure 4).

**Figure 4:**
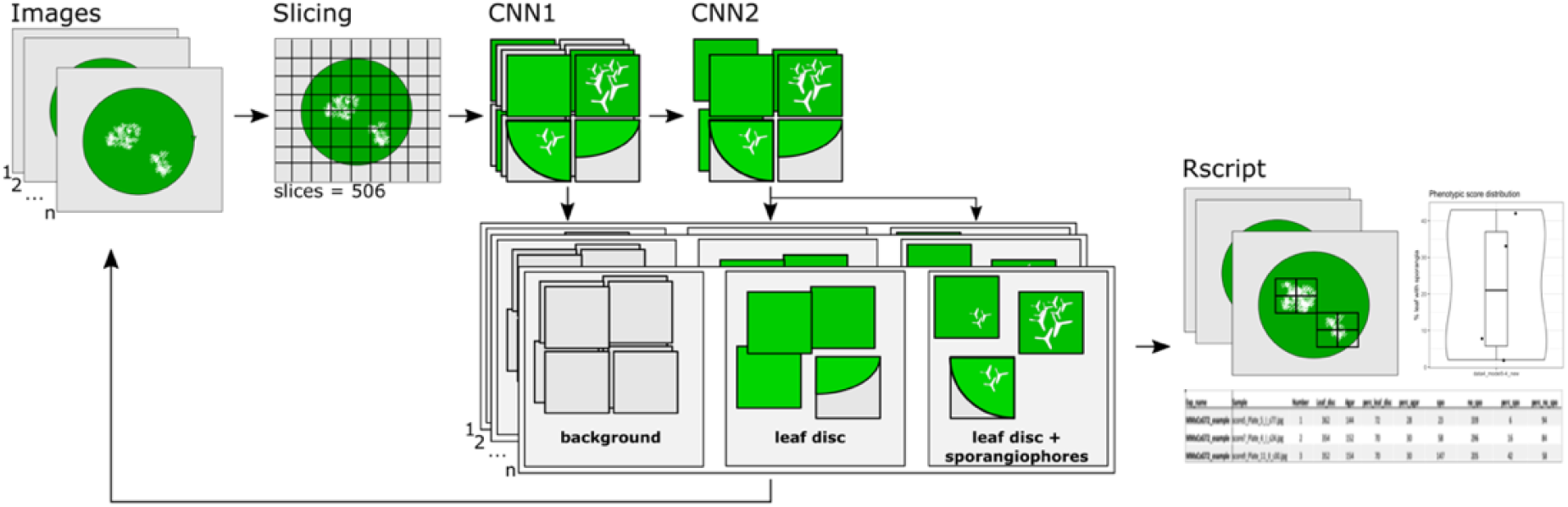
Scheme of the classification process of the two neuronal-networks.

First, a leaf disc image is sliced into 506 pieces which are then classified in water agar (background) or leaf disc. Leaf disc images are further analyzed with the second CNN2, which classifies them into leaf disc without and leaf disc with sporangiophores (Figure 4). The pipeline loops over all images in a supplied folder and after classification of all image slices of all images a R script plots the location of sporangiophores on the original RGB image, generates a plot with the percentage of leaf disc area with sporangiophores and a table of all generated values (Figure 4). As control, all images of classified sporangiophores are stored in a folder for manual inspection. The pipeline together with detailed installation instructions is available under www.github.com/Daniel-Ze/Leaf-disc-scoring. Additional information on how the CNNs were trained are included in the repository in the sub-section “scripts”.

## 3. Results

### 3.1 SCNN training results

The results of model training with the two SCNNs are shown in Figure 5. The first SCNN (background vs. leaf disc; CNN1) achieved an overall validation accuracy of 98% and a validation loss of only 6%.

**Figure 5:**
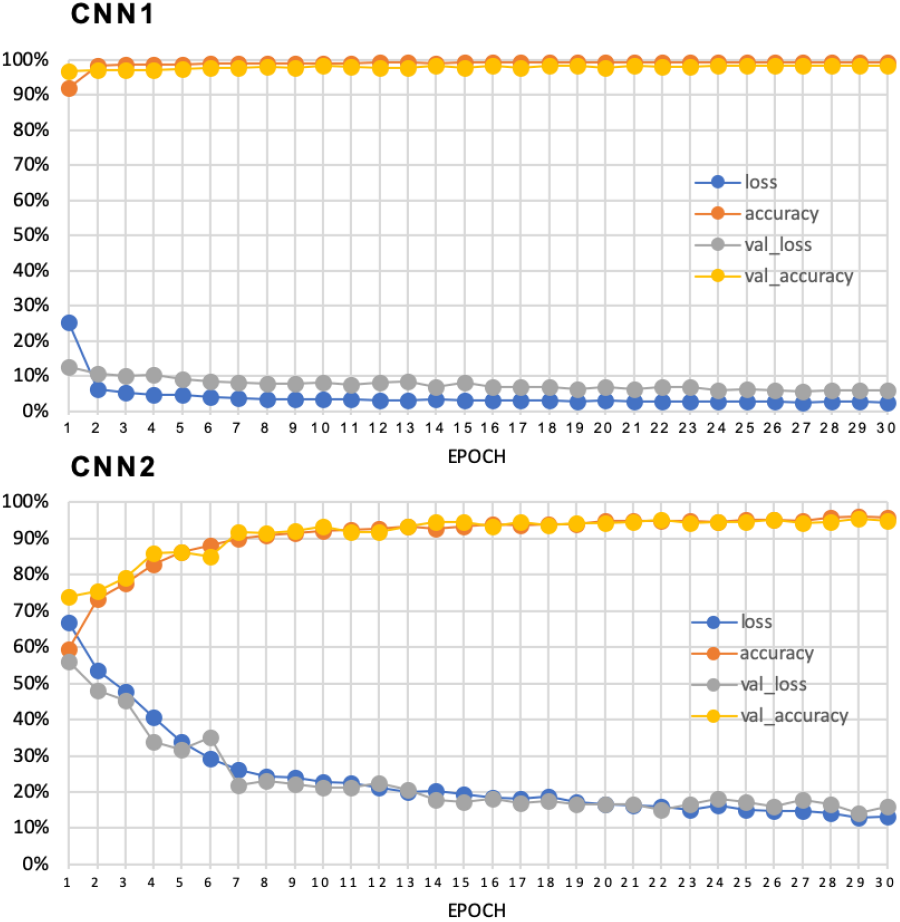
Training of the two SCNNs for background vs leaf disc (CNN1) and sporangiophores vs no sporangiophores (CNN2). Training / validation accuracy and loss (binary crossentropy) are plotted for 30 epochs (loss = training loss, val_loss = validation loss, accuracy = training accuracy, val_accuracy = validation accuracy).

This model was supposed to be the easier one of the two SCNNs as the image slices of background and the leaf disc itself differed the most (Figure 5). The second CNN (sporangiophores vs. no sporangiophores; CNN2) achieved a slightly lower accuracy. Overall validation accuracy after 30 epochs of training of this model was at 95%. The validation loss was higher with around 15% (Figure 5). The models were in both cases fitting as shown by the overlay of the training and validation curves (Figure 5). Both results are very satisfying and indicate a perfect basis for predicting the different classes in the leaf disc scoring pipeline. Code and image data to reproduce the results are available in the supplied GitHub repository.

### 3.2 Ground truth data

In total, 30 leaf discs were classified manually. The leaf disc set is composed of 15 leaf discs from the cross of ‘Morio Muskat’ x COxGT2 and 15 from the cross of ‘Cabernet Dorsa’ x Couderc 13. The original RGB picture together with the ground truth data from three independently scoring experts is plotted together with the CNN predicted image slice classes (Figure 6).

**Figure 6:**
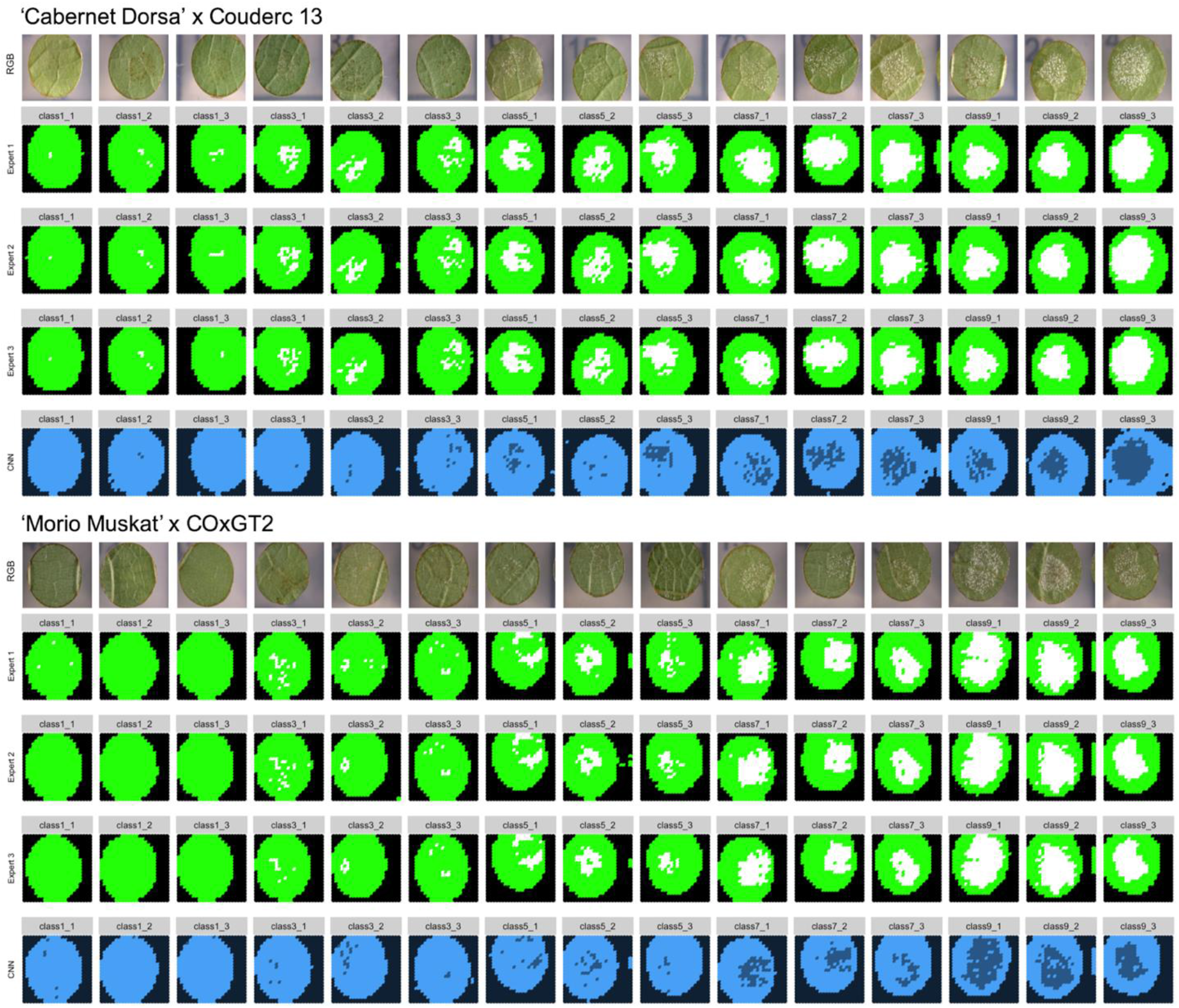
Original leaf disc pictures from the two different genetic backgrounds together with ground truth data and the results of the SCNN classification of the image slices. (black = background (1), dark blue = sporangiophores (2), light blue = no sporangiophores (3))

The ground truth data generated independently by the three experts are visually in a very high accordance with the DM infections on the leaf disc RGB images. Furthermore, all three manual evaluations of the experts show high similarity to each other. The results of the leaf disc scoring SCNN pipeline is plotted below the generated ground truth data of the three experts in Figure 6. The background and leaf disc itself are classified very consistently. However, there are a few discrepancies for the images from the unrelated genetic background. For example, in the image class7_3 we see a problem due to a partial second leaf disc in the picture. Besides this the classification of the first CNN is working very well. Looking at the image slices grouped into “leaf with sporangiophores” it shows that these are always predicted in the right areas of the leaf disc compared to the RGB picture and the ground truth data (Figure 6). However, apparently the second CNN is not detecting all image slices with sporangiophores. This issue seems to be a bit more pronounced for the leaf disc images from the ‘Cabernet Dorsa’ x Couderc 13 cross which was not used for training of the neural network (Figure 6).

### 3.3 Performance evaluation of the neural network

As an objective evaluation of the SCNN the manually assigned classes shown in Figure 7 were compared to the predicted SCNN classes. From that we calculated the percentage of true positives for each leaf disc image consisting of 506 sub-images.

**Figure 7:**
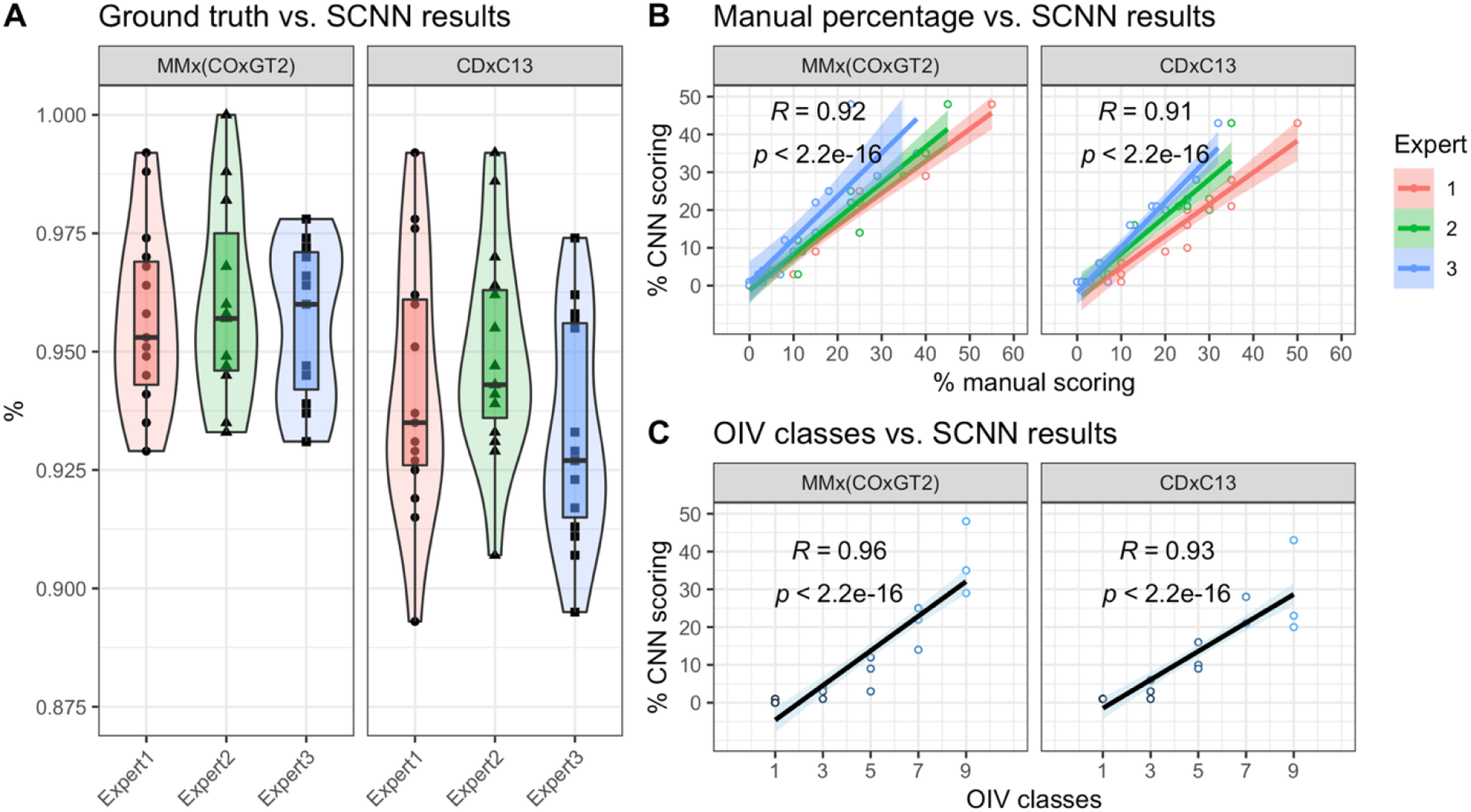
**A** True positive classifications of the CNNs compared to independent manual classification of three experts. **B** Manually scored percentage leaf disc area with sporangiophores correlated with the CNN percentage output. The confidence intervals for the linear regressions are indicated together with the overall correlation coefficient. **C** Assigned OIV classes of the leaf discs correlated with the percentage leaf disc area covered with sporangiophores calculated by the SCNN. (MMx(COxGT2) (‘Morio Muskat’ x COxGT2) = population used for SCNN training; CDxC13 (‘Cabernet Dorsa’ x Couderc 13) = population used for validation)

The results are shown in Figure 7A. The median percentage of true positive classifications ranges for the 15 ‘Morio Muskat’ x COxGT2 derived leaf disc images from 96 – 97%. The overall percentage of true positives ranges from ∼92 – 100% (Figure 7A, MMx(COxGT2)). The median CNN classification of true positive values ranged between 92 – 94% for the untrained ‘Cabernet Dorsa’ x Couderc 13 genetic background. The overall range of true positives was between ∼ 89 – 99% (Figure 7A, CDxC13).

Classifying 506 image slices for each leaf disc by hand is far from a real-world application. Therefore, the three experts manually estimated the percentage leaf disc area covered with sporangiophores and the inverse OIV452-1 classes available for each leaf disc were used to check the performance of the SCNN (Figure 7B and C).

The leaf disc scoring pipeline automatically calculates the percentage leaf disc area covered with sporangiophores which made it easy to correlate the output with the manually estimated percentages. For the leaf disc images from the cross ‘Morio Muskat’ x COxGT2 the correlation coefficient was 0.92 with a p-value well below 0.001 (Figure 7B, MMx(COxGT2)). For the ‘Cabernet Dorsa’ x Couderc 13 leaf disc images the correlation coefficient was with 0.91 a bit lower but nonetheless highly significant with a p-value also well below 0.001 (Figure 7B, CDxC13). The linear regressions of the three independent experts overlay partially for both datasets. This overlap is higher in lower percentages and decreases with higher estimated percentages (Figure 7B).

The final correlation was performed with the percentage of leaf disc area with sporangiophores from the leaf disc scoring pipeline and the assigned OIV452-1 classes. All three experts assigned the same classes for the different leaf disc images, therefore only one linear regression is indicated. The correlation coefficient for the cross ‘Morio Muskat’ x COxGT2 is again at 0.96. For pictures from the ‘Cabernet Dorsa’ x Couderc 13 cross the correlation coefficient is 0.92. For both correlations, the p-value is well below 0.001 (Figure 7C). The results for true positives and the two different correlations suggest that the output of the CNN classification is fitting very well with the values that the independent experts have manually estimated independently for the different datasets.

### 3.4 Leaf disc scoring pipeline

As mentioned before, the whole scoring process was combined into a single pipeline that can be executed via the command line in a Unix environment (Figure 4).

To visualize the leaf disc scoring pipeline results, we implemented a R script that automatically plots the positions of the predicted image slices with sporangiophores onto the original RGB image (Figure 8A). The R script further produces a plot with the phenotypic score distribution according to the observed percentage leaf disc area covered with sporangiophores (Figure 8B). Together with this a tab delimited table is generated storing the gained information necessary for downstream analysis of the data (Figure 8C). We made the whole pipeline publicly available on GitHub as an open source repository (https://www.github.com/Daniel-Ze/Leaf-disc-scoring). Along with the pipeline, we included a detailed walk-through on how to train simple but efficient SCNNs with little image data on small scale hardware like laptops or desktop PCs without the need of a dedicated high-performance GPU. The Jupyter notebook included in the repository gives detailed instructions on how the SCNNs were trained and how the SCNNs can be trained for other traits or other plants as long as a leaf disc scoring is possible. The pictures used to train the two binary SCNNs are included as datasets in the repository. Together with the Jupyter notebook they can be used to track the process.

**Figure 8:**
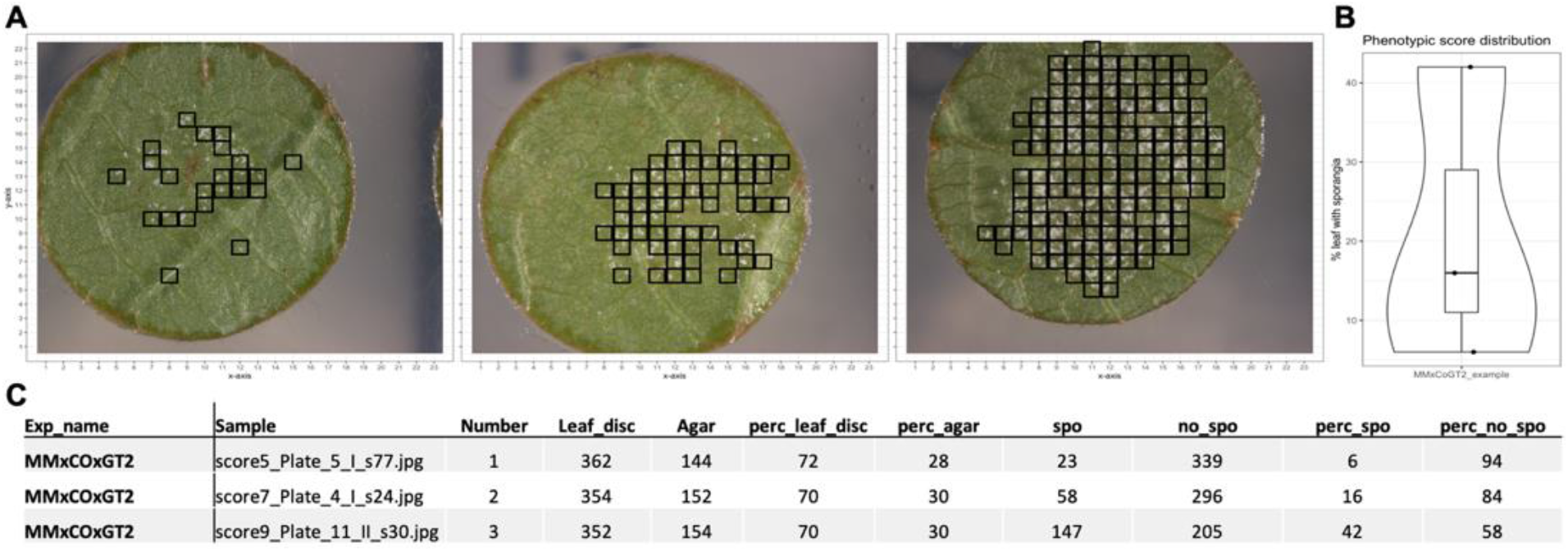
Example results from the leaf disc scoring pipeline. **A** Original RGB images with indicated leaf disc slices classified as “with sporangiophores”. **B** Overview of calculated percentage leaf disc areas for the analyzed set of images. **C** Tab delimited table output of the leaf disc scoring pipeline that can easily be integrated in downstream analyses. .(Exp_name –experiment name, Sample – Sample id used in the analysis, Number-serial number, Leaf_disc- slices identified as leaf, Agar- slices identified as agar/background, Spo- slices identified as sporangiosphores infected, perc – percentage)

## 4. Discussion

We present here an automated leaf disc scoring pipeline using RGB images and trained shallow convolutional neural networks (SCNNs). We further supply detailed instructions on how to train these SCNNs for novel traits to adjust the pipeline accordingly. As proof of concept, we trained the neural networks to efficiently detect grapevine downy mildew emerging from inoculated leaf discs from two segregating populations generated from different genetic backgrounds. A similar approach was already successfully implemented for the detection of grapevine powdery mildew caused by *Erysiphe necator* [24]. The pipeline presented here allows the objective high-throughput screening of leaf discs with controlled infections, which is a crucial part of any plant breeding or QTL mapping workflow. In recent years, the use of deep learning for image classification in the area of plant pathogen detection has received much attention. For reviews on this topic see [6,7,28,29]. For visually easily detectable diseases located on the leaf surface like powdery mildew several deep learning image segmentation approaches have been published in recent years [24,30,31].

As downy mildew is a major problem in temperate climates, where relatively cold and humid spring times are common, a major effort is made to breed new grapevine cultivars with resistance against this pathogen. As for all breeding programs objective phenotyping of the offspring of a segregating cross is essential. This requires highly trained personnel who have many years of experience with the pathogen of choice. However, in the meantime, breeding programs are forced to increase their capacity and the need for high-throughput phenotyping methods emerges. Leaf disc assays have already been an important part of resistance scoring under controlled laboratory conditions for a long time. The evaluation of these tests however, was done manually in most of the cases [32–34]. The advancements in computer vision and machine learning in recent years enabled a mainstream application of previously highly complex and very computationally intense algorithms. The well documented open source projects TensorFlow [2] and Keras [26] built on top of the latter have made it easy for people with limited knowledge in Python programming to apply these principles to their own datasets.

The pipeline downstream of the image acquisition presented in this study was designed as open source repository for anybody to be used and can be adjusted to any individual needs. We assume that image acquisition will differ from laboratory to laboratory therefore we put together a workflow for training new SCNN models with image data available from other image acquisition pipelines. However, we also included detailed instructions and a template for automatic imaging with a Zeiss Axio Zoom v16 with ZenBlue3.0 software.

### 4.1 SCNN versus expert evaluation

The goal of this study was to create an efficient method for scoring disease severity on leaf discs using a limited amount of image data and a limited amount of computational power. To achieve this, we used simple SCNNs and trained these with small numbers of images. Further, we decided to use only a simple binary classification of the images reducing the complexity and allowing to first distinguish background from leaf disc and then, in the second step the detection of infected leaf disc areas. This approach will allow specific training of the second SCNN for additional traits or pathogens therefore increasing the modularity of the pipeline.

Training of the first SCNN shows that the detection of leaf disc image slices is a fairly easy task for the neural network as the leaf slices differ greatly from the background. Training of this SCNN showed a high specificity with a 98% validation accuracy and a validation loss below 10% (Figure 5). The training of the second SCNN was thought to be much more complicated in terms of distinguishing leaf hairs from sporangiophores. However, after adjustments to the model such as increasing the number of image convolutions together with two additional fully connected layers and a different model optimizer, training of this neural network also showed a very high validation accuracy of 95% with a validation loss around 15% (Figure 5).

To evaluate the performance of these trained SCNNs, three independent experts sorted 15180 image slices (30 leaf discs) into the three categories: background, leaf disc with and leaf disc without sporangiophores (Figure 2). A feature of trained SCNNs is their transferability, meaning that in optimal conditions the SCNNs should have learned distinct features allowing a categorization of images previously not observed by the SCNN. To test this, we chose 15 new images from the same genetic background plus 15 images from another cross with an unrelated genetic background as testing set. The test set of images was analyzed with the two SCNNs and the ratio of true positives was calculated from that as means of performance evaluation. The median ratio of true positives was around 96% for the trained genetic background and 93% for the unrelated genetic background. This indicates that the transferability of the two SCNNs is given with only a slightly decreased accuracy (Figure 7A).

Correlating the results of the two SCNNs with real world applications such as manual estimation of sporangiophores-covered area and OIV452-1 classes showed highly significant correlations again with a slight decrease for the unrelated genetic background. The subjectivity of the manually estimated area covered with sporangiophores is indicated by the limited overlap of the three linear regressions (Figure 7B). This was not the case for the assigned inverse OIV452-1 classes due to the reduction of complexity by the limited number of assignable classes. For these two correlations again, a slight decrease was observed for the second unrelated genetic background as it was seen for the true positive ratio (Figure 7C).

To counteract this observed difference, we propose to generate a diverse training set composed of multiple different genetic backgrounds to maximize the transferability and increase the already very good performance of the two SCNNs. The results of the true positive ratio and the performed correlations indicate that the results of the two SCNNs are a well-suited objective substitute for these two subjective scoring methods.

### 4.2 The leaf disc scoring pipeline

We demonstrated that the two SCNNs can efficiently detect grapevine DM infections on images of leaf discs. This allows their incorporation as a valuable tool for the evaluation of resistance traits in breeding processes and for the characterization of genetic resources in germplasm repositories. The new leaf disc scoring method is particularly applicable for specific research purposes like the phenotyping of large populations needed for the detection and QTL mapping of novel resistances to e.g. DM. Therefore, we wrote a simple open-source pipeline incorporating the two CNNs. The pipeline is an easy to execute command-line application that was tested to work on Unix systems.

With the modular two-step detection process, it will be easy to train the second CNN to detect and quantify other grapevine pathogens, where leaf infections are currently only visually assed like *Guignardia bidwellii* causing black rot, *Colletotrichum gloeosporioides* causing ripe rot, *Phakopsora euvitis* causing leaf rust or *Elsinoë ampelina* causing anthracnose [35–38]. Grapevine DM is by far one of the easiest pathogens to be detected with its very white sporangiophores. For other pathogens such as *E. necator* the used SCNN model layers of CNN2 most probably have to be adjusted to a DCNN like in Biermann et al. [24] to fit the different fungal structures formed during the infection. However, we are confident that this scoring pipeline will find a multitude of applications, not only for resistance assessment of grapevine diseases, but also for other plant-pathogen systems where leaf disc assays are already established or will be developed in the future. The whole process for training the SCNNs is documented in an easy to use Jupyter notebook (see GitHub repository).

### 4.3 Further future improvements

A step in improving the leaf disc scoring pipeline would be to switch from binary to categorical classification. This would allow the detection of several different traits with the second CNN at the same time. An even better way of categorical classification would be semantic labeling of the traits analyzed on the leaf disc image slices. This would allow a much better accuracy in terms of detection. For example, the investigated leaf discs infected with DM showed different sporangiophore densities. Such differences are neglected by the CNNs presented here. To further improve the detection accuracy a switch to alternative deep learning techniques such as semantic labelling is inevitable. In a recent study this approach was demonstrated for black rot, black measles, leaf blight and mites on grapevine leaves [39]. Further studies have shown promising detection of leaf localized diseases using semantic segmentation for coffee and rice [40,41] However, the results obtained with the simple binary classification setup presented here clearly show a high correlation with standard scoring methods indicating its suitability for high-throughput phenotyping of leaf discs. Furthermore, it keeps the requirements in terms of computational resources and image training sets to a minimum allowing a fast adaptation to new plant-pathogen systems.

## 5. Conclusions

The pipeline itself for image classification is a useful tool for automated analysis of image data. This process is usually carried out by highly trained personnel that have long-term experience with the investigated pathogen and disease. One, or for improved objectivity, two experts have to score the inoculated samples manually to obtain the most objective classification results. With the pipeline presented here most time is spent on preparing the inoculation experiment and subsequently recording the image data. This scoring pipeline removes the subjective manual scoring. It will ensure consistent scoring results throughout all experiments. Additionally, the whole pipeline can be transferred and adjusted to new pathogen-plant systems with a minimal set of training images. It is a valuable tool in different agronomic fields from basic research to plant breeding.

## Author Contributions

D.Z. designed and supervised the experiment, designed and trained the neural networks, wrote the program code, performed the statistical analysis, generated the ground truth data, wrote initial draft. N.M. performed artificial inoculations, recorded image and phenotypic data, classified the training data, generated the ground truth data, helped writing the initial draft. A.S. performed artificial inoculations, recorded image data, generated the ground truth data, helped writing initial draft. R. T. supplied experimental infrastructure and reviewed and edited the manuscript. R.T., L.H. and E.Z. promoted this work, provided funding, reviewed and edited the manuscript.

## Funding

This research was funded by DFG (Zy11/9-1 and 9-2) and by BMEL & BÖLN (VitiFit).

## Data Availability Statement

The data for the presented SCNNs including scripts and images are available in the corresponding GitHub repository: www.github.com/Daniel-Ze/Leaf-disc-scoring

## Conflicts of Interest

The authors declare no conflict of interest. The funders had no role in the design of the study; in the collection, analyses, or interpretation of data; in the writing of the manuscript; or in the decision to publish the results.

## Notes

https://github.com/Daniel-Ze/Leaf-disc-scoring

### Competing Interest Statement

The authors have declared no competing interest.

